# CGPA: multi-context insights from the cancer gene prognosis atlas

**DOI:** 10.1101/2024.07.19.604345

**Authors:** Biwei Cao, Xiaoqing Yu, Gullermo Gonzalez, Amith R Murthy, Tingyi Li, Yuanyuan Shen, Sijie Yao, Jose R. Conejo-Garcia, Peng Jiang, Xuefeng Wang

## Abstract

Cancer transcriptomic data are used extensively to interrogate the prognostic value of targeted genes, yet basic scientists and clinicians have predominantly relied on univariable survival analysis for this purpose. This method often fails to capture the full prognostic potential and contextual relevance of the genes under study, inadvertently omitting a group of genes we term univariable missed-opportunity prognostic (UMOP) genes. Recognizing the complexity of revealing multifaceted prognostic implications, especially when extending the analysis to include various covariates and thresholds, we present the Cancer Gene Prognosis Atlas (CGPA). This platform greatly enhances gene-centric biomarker research across cancer types by offering an interactive and user-friendly interface for highly customized, in-depth prognostic analysis. CGPA notably supports data-driven exploration of gene pairs and gene-hallmark relationships, elucidating key composite biological mechanisms like synthetic lethality and immunosuppression. It further expands its capabilities to assess multi-gene panels using both public and user-provided data, facilitating a seamless mechanism-to-machine analysis. Additionally, CGPA features a designated portal for discovering prognostic gene modules using curated cancer immunotherapy data. Ultimately, CGPA’s comprehensive, accessible tools allow cancer researchers, including those without statistical expertise, to precisely investigate the prognostic landscape of genes, customizing the model to fit specific research hypotheses and enhancing biomarker discovery and validation through a synergy of mechanistic and data-driven strategies.

**Significance:** CGPA is a streamlined, interactive platform for multi-context gene-centric prognostic analysis, simplifying biomarker discovery and validation in oncology for clinicians and basic scientists, and bridging a critical gap in translational cancer research.

## INTRODUCTION

In the cancer genomics field, the transcriptome-wide RNA-seq approach is one of the dominant approaches for molecular profiling of tumor tissues. Over the past decade, an exponential growth in cancer-related RNA-seq data has been witnessed, much of which has been deposited into public domain. Due to its widely expanded utilization and accessibility, tumor transcriptome databases have become indispensable resources for both preclinical research and clinical testing. Despite the rapidly growing data in repositories such as GEO and SRA, the Cancer Genome Atlas (TCGA) remains the largest and most commonly used pan-cancer database, which contains curated transcriptome data from over 11,000 primary cancer samples across 33 major cancer types^1,2^. However, the official portal for accessing TCGA data is not user-friendly enough for users without sufficient programming and data manipulation skills. For example, in the Genomic Data Commons (GDC) Data Portal, the Level-3 RNA-seq-based expression (e.g. RPKM) data are scattered across many separate files store information per patient sample. Therefore, the online analysis website cBioPortal^3^ has been a more popular choice, and a de facto go-to resource, among clinicians and basic science researchers, especially for initial data mining. Although cBioPortal provides very comprehensive gene and mutation spectrum analysis across patients, it has limited functions to explore the association between gene expression (GE) and clinical outcomes such as patient survival. Another online database, GEPIA^4^, offers more advanced correlation analysis of GE data and allows for a more straightforward gene-based query based on data curated from TCGA and GTEx. Unfortunately, due to the location of its web server, access to this website is not universally granted by institutions or entities in countries/regions such as the United States.

This work has been motivated by several observations in oncological research. First and foremost, the analysis of continuous gene expression data presents distinct challenges when compared to the binary mutational data. The prognostic modelling of gene expression data is inherently sensitive to the selection of cutoffs. A significant limitation in current tools is their reliance on fixed cutoffs, most commonly the median and quartiles, without the flexibility to adjust these thresholds based on specific study needs. Determining an optimal threshold for these expression levels, to classify patients into different risk groups, can significantly impact study outcomes, potentially leading to varying conclusions even with identical datasets. Yet, many published studies, especially in basic science settings, have only shown the univariate association between survival and GE level without adjusting for any clinical outcomes. This is most commonly visualized by the separation of Kaplan Meier (KM) curves from high- and low-GE patient groups. Such practices can inadvertently yield inaccurate or even misleading conclusions, especially considering the prognostic association of many genes can be confounded by factors such as sex, known cancer subtypes, tumor grade and stage.

A second key motivation behind our work stems from arises from the observed limitations of current tools in constructing comprehensive multi-gene prognostic signatures. Many investigators are transitioning from genome-wide searches to more hypothesis-driven approaches. Driven by prior knowledge and reinforced by preliminary findings, researchers are increasingly focusing on specific gene pathways. They postulate that these curated pathways may harbor significant prognostic insights, potentially serving as key determinants in understanding disease progression and patient outcomes. Moreover, as translational studies progress, there is an increasing trend towards developing gene panels that encompass a curated set of genes. These panels aim to provide a more holistic view of the disease, capturing the collective influence of multiple genes. However, the development and validation of such panels require sophisticated analytical tools that can cohesively evaluate the joint impact of all included genes. Finally, an extremely important but often ignored fact is that most existing cancer GE data were generated from bulk tumor tissue and gene expression are highly sensitive to cellular composition in the sampled tissue blocks. And thus, GE values can be strongly influenced by the content of cancerous cells as well as the immune cell infiltration in the tumor microenvironment (TME). Accordingly, many top survival-associated genes, especially in immunogenic cancer types, are less ‘functional’ themselves but merely caused by their strong correlation with tumor purity and TME scores. We reason that this issue can be alleviated by considering multiple genes and molecular scores simultaneously in the survival model. Following this multi-gene perspective, there is also an emerging recognition of the importance of two-gene, or gene-pair, analyses. Such an approach, while seemingly simpler than multi-gene analyses, holds its own set of complexities and advantages. Gene-pair analyses also empower researchers to discern critical gene-gene interactions, shedding light on synergistic or antagonistic relationships that can be instrumental in shaping cancer progression and patient outcomes. A prime example is synthetic lethality, which arises when a combination of deficiencies in the expression of two genes results in cell death, potentially leading to improved patient survival.

In response to above-discussed problems, we introduce CGPA (Cancer Gene Prognosis Atlas). This innovative online tool and interactive analysis portal is meticulously designed to address the complex needs of gene-centric biomarker discovery and validation. CGPA provides fast access and intuitive query to prognostic significance of a gene signature across a pan-cancer spectrum. The tool fills a gap in the community as the first interactive tool for generating publication-ready results from multivariable and multi-gene survival models. CGAP contains several innovative functions that allows effective mining of correlation structure among targeted genes, as well as the function to generate and test the pooled GE signatures based on customizable gene panels. Taken together, CGPA is a timely and unique resource that will complement the existing databases in cancer genomics, and will immediately facilitate the process of gene validation, prognostic signature discovery, and exploration of therapeutic targets.

## RESULTS

### CGPA is a proactive and interactive tool for gene-centric prognostic analysis

The development of the CGPA platform was driven by the observation that many published studies depend solely on univariable Kaplan-Meier plots for the exploration and validation of the prognostic relevance of specific genes. Within this context, GraphPad Prism emerges as the most popular software for such analysis among clinicians and non-statisticians in basic science laboratories. One common misconception is that multivariable analysis, by introducing more covariates and thus increased degrees of freedom, might reduce statistical power and render findings of marginal significance to less or non-significant outcomes. Contrary to this belief, CGPA’s proactive multivariable analysis, by adjusting for essential covariates, has identified genes whose significance becomes more apparent, termed here as univariable missed-opportunity prognostic (UMOP) genes. Building on this foundational concept, the CGPA has evolved to include a suite of innovative functions specifically designed to enhance gene-centric prognostic analysis. Among its unique features, we highlight a few that distinguish it from existing databases: (1) CGPA provides a quick and holistic overview of the pan-cancer prognostic landscape, allowing users to assess the prognostic value of genes across different cancers. (2) It offers detailed, customized gene-based survival analyses, including flexible cutoffs in Kaplan-Meier plots, covariate-adjusted Kaplan-Meier, and multivariable Cox models, accommodating diverse research needs. (3) It provides a designated tab called “ProgSplicing” to further explore the prognostic pan-cancer landscape of alternative splicing events. (4) The platform supports in-depth exploration of gene pairs and gene-hallmark interactions, shedding light on complex mechanisms like synthetic lethality and immunosuppression that are pivotal to cancer biology. (5) Additionally, CGPA extends its capabilities to the evaluation of multi-gene panels through a mechanism-to-machine approach. It also provides an option for users to analyze gene signatures by uploading their own data set. Following an extensive two-year period of internal testing at Moffitt, CGPA’s public web portal has received more than 194,000 visits by March 26, 2024.

As illustrated in **Figure 1**, the CGPA web application have three main modules. At the forefront is its comprehensive single-gene prognostic discovery. While many studies rely on univariate regression or Kaplan-Meier plots for their interpretative simplicity, such methods can often lead to biased insights. Factors like tumor purity, patient heterogeneity and hidden confounding variables might distort the association results. However, incorporating additional factors into the model can lead to a reduction in statistical power, a consequence of the increased degrees of freedom. Therefore, both univariable and multivariable analysis should be considered for a more holistic and robust understanding of gene prognostic significance. Second, CGPA introduces the gene-pair interaction model, which models two genes simultaneously. The rationale for examining gene interactions is rooted in the understanding that genes often don’t function alone; their combined effects can play pivotal roles in cellular pathways and responses. By focusing on pairs of genes, CGPA offers researchers to test the prognosis in a specific context. For example, when an investigator has a specific target gene in mind, analyzing its interaction with another gene might offer additional context. The two-gene model can also help in exploring potential synergistic effects, which can sometimes be more straightforward to interpret than interactions spanning multiple genes. Such interactions can shed light on mechanisms like synthetic lethality and synthetic viability. While several tools in the field offer gene interaction analyses, many might not probe into the granularity of two-gene interactions or might have a more specific focus, such as TIDE’s emphasis on immune signatures^5^. Lastly, CGPA’s multi-gene panel discovery feature emerges as a response to the complexities inherent in cancer genomics. Gene pathways often comprise a vast array of genes, and due to the inherent heterogeneity across cancer types, the prognostic significance of these genes can sometimes become diluted when tested as a combined score. Recognizing this challenge, CGPA goes beyond just examining gene enrichment scores and prognostic signatures. It goes further by segmenting larger gene groups into more manageable and biologically relevant subsets through subnetwork analysis. This breakdown not only allows for an investigation into the distinct roles of each gene subset but also offers insights into how these subsets might interact with one another. This strategy might help in revealing complex gene interactions or pinpointing gene subsets linked to specific cancer traits. In summary, with its multi-tiered approach to gene analysis, CGPA serves as a holistic gene-based tool, aiming to fill the gaps in cancer genomics research and provide a thorough analysis for both basic and translational studies.

**Figure 1.**
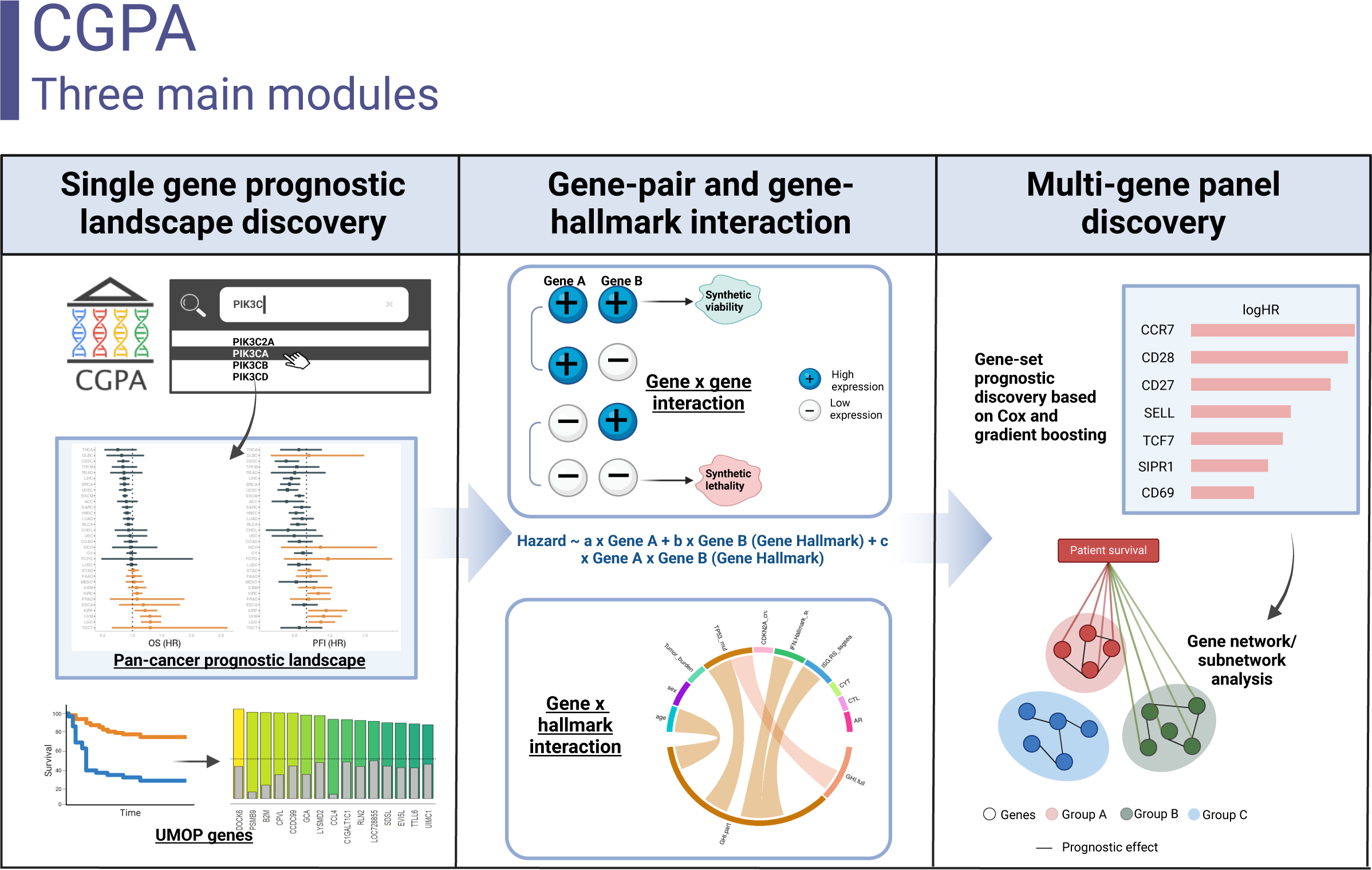
Overview of main functions in CGPA to facilitate gene-centric prognostic biomarker discovery and validation.

### Single-gene module and univariable missed-opportunity prognostic (UMOP) genes

The overarching goal of the single-gene module is to provide a comprehensive evaluation of gene prognostic significance, focusing on critical aspects of biomarker validation that existing tools have not yet addressed. When a gene is queried in the search bar, CGPA first renders a prognostic summary across various cancer types. The prognostic significance of the gene is illustrated based on the forest plot ranked by the estimated effect size on survival outcomes, e.g., hazard ratio (HR). This visualization method presents a pan-cancer overview of how the gene’s expression influences patient OS and PFI. Each line in the forest plot represents a cancer type, with the position and length of the line indicating the estimated HR and its confidence interval, respectively. A line crossing the vertical line of no effect (hazard ratio of 1) suggests that the gene does not significantly impact survival, whereas lines to the left or right indicate favorable or adverse effects, respectively. As an ad-hoc exploration, those cancer types that demonstrate significant prognostic impact will be further explored using the Kaplan-Meier plot by groups stratified based on optimal cutoff points. These cutoff points are identified through Maximally Selected Rank Statistics^6,7^, a method that selects the optimal threshold maximizing the difference in survival metrics between high and low expression groups. However, this data-driven approach carries a risk of overfitting, and researchers should be cautious when the stratified group is very unbalanced, or the direction of the risk is inconsistent within the HR estimate. To elucidate the biological context of the genes, the prognostic summary panel also incorporates figures that display cancer-level gene expression profiles in tumor and normal tissues, alongside the protein-protein interaction network based on the STRING database. The more customized and detailed examination of gene-based analysis is available when click the “multivariable analysis” tab, where users can further explore the impact of a gene on patient survival within the context of clinical covariates and tumor purity. To facilitate visualization, this tab provides results from Covariate-adjusted KM functions^8^ alongside results from multivariable Cox regression analysis. In the adjusted KM plots, users have the flexibility to choose between median, quartile, or optimal-cutoff based stratification to visualize survival curves and explore clinical disparities.

In our comprehensive analysis of pan-cancer transcriptome data, we have identified a multitude of genes with prognostic potential that have consistently been missed by a univariable Cox regression analysis. Below, we share insights based on results from a group of immunogenic cancers, including bladder cancer (BLCA), head and neck squamous cell carcinoma (HNSC), lung adenocarcinoma (LUAD), skin cutaneous melanoma (SKCM), and uterine corpus endometrial carcinoma (UCEC). We directed our attention to those genes that exhibited prognostic significance in the multivariable model but not when analyzed univariately, suggesting their potential independent influence beyond clinical covariates and general immune scores. We refer to them as univariable missed-opportunity prognostic (UMOP) genes. **Figure 2** illustrates this contrast with bar plots that compare the -log10 p-values from both a univariable Cox model and a multivariable Cox model that adjusts for age, sex, tumor purity, and cytotoxic T lymphocyte (CTL) levels. In BLCA, our analysis identified a diverse set of genes as UMOP biomarkers. These genes are implicated in a variety of cellular or immune functions including extracellular matrix remodeling (MMP15, HPSE), metabolic alternations (PDXK, BCAT2), immune response (C1QB, C1QC), signal transduction (MAP3K7, CAMK4), and critical gene regulation (MYCL1, ZNF436). In HNSC, the top UMOP genes that were missed by the univariable model include DOCK6, PSMB9, B2M, CPVL, and CCDC99. Remarkably, a set of well-known immune-related genes such as HLA-C, HLA-H, HLA-B, and HLA-F, also demonstrated prognostic significance in the multivariable analysis. This observation reinforces the notion that certain immune gene signatures may confer independent prognostic value, distinguishing themselves from the broader correlations typically seen in immune gene expression within the context of immunogenic cancers. Furthermore, genes with key roles in immune responses, such as IL15, IFNB1, and SERPINA1, have also been recognized as UMOP genes in our multivariable analyses, adding a new layer to our understanding of the complex interactions in the immune landscape of head and neck cancer. In LUAD, our analysis also uncovered a set of immune modulator genes such as TAP1 and GBP1, alongside the interferon signaling mediator STAT1, suggesting a significant role of immune surveillance and response in LUAD progression. In addition, we identified several genes integral to DNA replication and repair processes, such as PCNA and RFWD3, which may indicate genomic instability as a prognostic indicator in lung cancer. The identification of SOD2, which regulates oxidative stress responses, and MT1X, a gene involved in metal ion homeostasis, highlights the balance between cellular defense mechanisms and metal regulation. In SKCM, our analysis intriguingly highlights HMOX1 and FTL, among the top UMOP genes, which are also involved in the cellular response to oxidative stress and iron homeostasis, respectively. Additionally, genes such as SIRPA, SIRPB1, VIPR1, ITLN1, and SELENBP1, are known to contribute to the interactions between cancer cells and the host’s immune system. These genes could be pivotal in modulating the effectiveness of both conventional therapies and immunotherapeutic approaches in skin cancer. In UCEC, our UMOP analysis further identified genes that are related to immune surveillance (e.g., MICB, CLEC2D, VCAM1, TNFA1P6, and TNFSF4) and genes that are related to cellular stress response (e.g., PPP2CA, GSTA2, HDAC8, and DNAJB6). Collectively, our comprehensive UMOP analysis in immunogenic cancers identified not only key immune genes, but also previously overlooked genes involved in other mechanisms that intersect with and influence immune-related processes.

**Figure 2.**
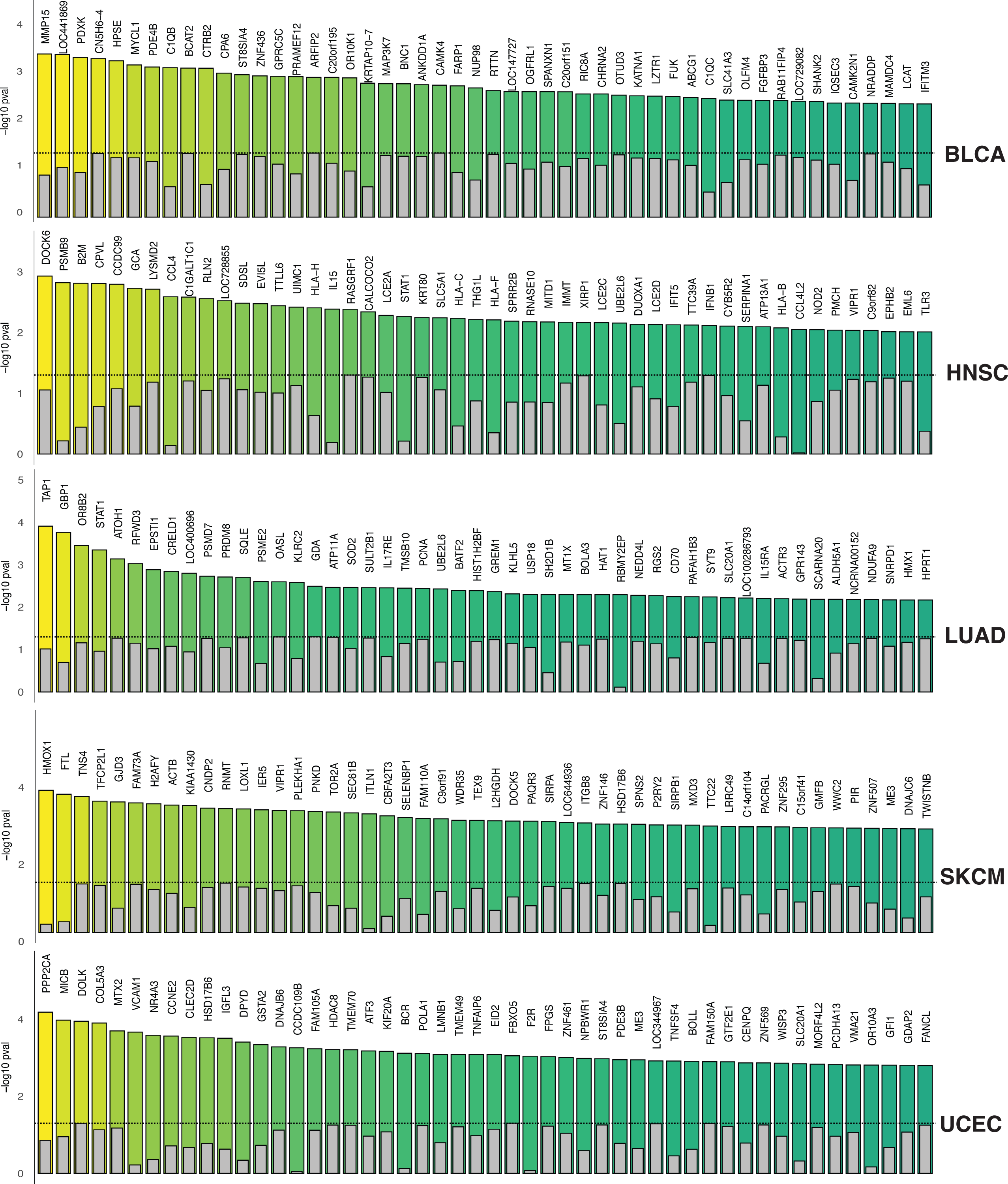
Identification of Univariable Missed-Opportunity Prognostic (UMOP) Genes across five immunogenic cancers: BLCA, HNSC, LUAD, SKCM and UCEC. Bar plots visualize the -log10 p-values from both univariable and multivariable analyses, highlighting the genes that exhibit significant prognostic potential in the multivariable context but are not reach statistical significance in the univariable analysis.

### Gene-hallmark interaction

CGPA offers an innovative feature for exploring gene-hallmark interactions through a dedicated “Gene-Hallmark Interaction” tab. This functionality allows users to investigate the interplay between targeted genes and cancer hallmark signature directly within a gene query. The investigation of gene-cancer hallmark interactions is rooted in the evolving understanding of cancer as not merely a collection of random genetic aberrations, but as a complex biological system characterized by specific, definable traits, i.e., hallmarks of cancer^9,10^. The hallmark covariates integrate a wide array of factors including top mutated genes, copy number variations (CNV), tumor mutational burden (TMB), immune scores (such as cytolytic activity (CYT)^11,12^ and cytotoxic T lymphocyte (CTL)^5^ levels), and the androgen receptor (AR) status, along with other hallmark signature scores. For each cancer type, we focused exclusively on the most prevalent mutations and copy number alterations (CNA) specific to that type; for instance, TP53 mutations and CDKN2A CNA in head and neck squamous cell carcinoma (HNSC). These scores are calculated based on gene expression data, utilizing single-sample Gene Set Enrichment Analysis (ssGSEA)^13^. Our investigation is based on the statistical framework, incorporating a multiplicative term in the Cox model to assess the interaction between two variables. We implemented two distinct models in this function tab: the Gene-Hallmark Interaction (GHI) full model and the GHI partial model. The full model evaluates the combined effect of genes and hallmarks, as well as their interactions, on patient survival (Hazard ∼ a.Gene + b.Hallmark + c.Gene x Hallmark), while the partial model only includes the main effect from the gene (Hazard ∼ a.Gene + c.Gene x Hallmark). This dual-model approach allows us to explore the potential interplay of genes and hallmarks under different assumptions. All interaction analysis allows for the adjustment of tumor purity in the analysis. The visualization of the interaction is facilitated through circos plots, in which significant interactions will be highlighted. When examining immune CTL scores as a hallmark, the model aligns with methodologies used in TIDE^5^ and ENLIGHT^14^, offering insights into immune evasion and response mechanisms by assessing the interaction’s impact on patient survival outcomes. **Figure 3** is an example of the pan-cancer gene-hallmark interactome of gene *IGSF8*, which is a novel innate immune checkpoint and a potential new target for immunotherapy.

**Figure 3.**
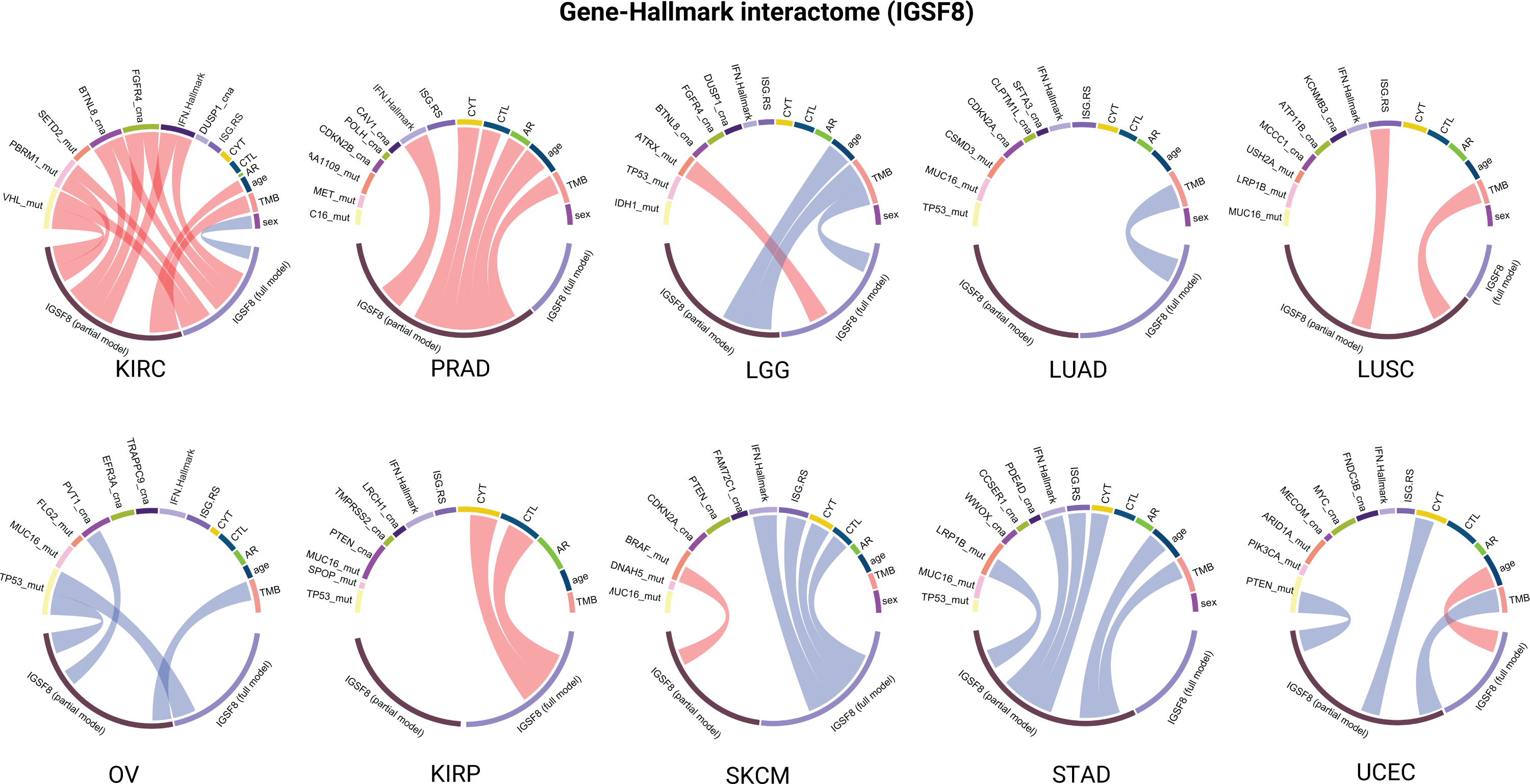
Pan-cancer gene-hallmark interactome of gene *IGSF8.* In the circos plot, significant interactions (at a significance level of 0.05) are represented as links. Positive interaction effects are indicated in red, while negative effects are shown in blue.

### Gene-pair interaction module

In this section, we briefly discuss the main results from utilizing the gene-pair interaction model within the CGPA toolkit, an important tool for understanding the prognostic value of gene interactions in cancer research. This module enables researchers to concurrently assess the survival impact of two genes, an important capability that none of the current cancer transcriptomic databases or tools offer. Users can click the “Two gene search” on the main page, which directs them to a dedicated analysis portal tailored for the in-depth prognostic analysis using the selected gene pair. As discussed previously, this module enables one gene anchored for a targeted or conditional survival modelling, facilitates interaction analysis crucial for identifying synthetic lethality and synthetic viability, and supports a more comprehensive multivariable model that accounts for interaction explicitly even without an interaction term^15^. It presents its findings through Kaplan-Meier (KM) plots and bivariable Cox regression analyses, the latter including interaction terms for a comprehensive evaluation. The simultaneous utilization of both KM and Cox models is essential, leveraging the straightforward, visual clarity of KM plots and the detailed, statistical insights provided by Cox models. While KM plots are easily interpretable, they can be sensitive to the choice of expression level cutoff. As demonstrated in **Figure 4**, the conventional approach to investigating gene interaction effects involves comparing patient survival across the four distinct quadrants created by dichotomizing gene expression levels into high and low categories. Figure 4A and 4B focus on the interaction between TCF7 and LAG3. TCF7, a transcription factor important for T-cell development and differentiation, and LAG3, an immune checkpoint that co-regulates T-cell activation, together highlight a complex regulatory mechanism that could be pivotal for immune evasion by tumors. Therefore, targeting the TCF7-LAG3 axis could improve existing immune checkpoint inhibitors by promoting a more robust and effective anti-tumor immune response^16^. In Figure 4A, patients with high TCF7 expression show a clear survival differentiation based on LAG3 levels, with high LAG3 levels being associated with better survival. In contrast, Figure 4B, focusing on the TCF7-low group, reveals no such survival distinction with varying LAG3 levels, indicating a significant interaction effect between TCF7 and LAG3. Figures 3C and 3D exemplify the concepts of synthetic viability and synthetic lethality, illustrating unique cases of gene-gene interaction where the survival outcomes of patients in one specific quadrant (high-high/low-low) are compared against those in the remaining three quadrants. For example, synthetic lethality arises when the simultaneous low expression or inactivation of two genes leads to cell death, whereas the low expression of either gene alone does not produce the same effect. Figure 4C illustrates synthetic viability using the VPS37A and DCLRE1B gene pair. Patients with high expression levels of both genes exhibit better survival compared to other stratified groups. Figure 4D presents a scenario of synthetic lethality with the UBE2Z and RNF117A gene pair. Here, patients characterized by low expression of both genes show improved survival outcomes compared to the other three groups. This finding suggests that therapies aimed at simultaneously inhibiting both genes could offer a strategic approach in treatment development.

**Figure 4.**
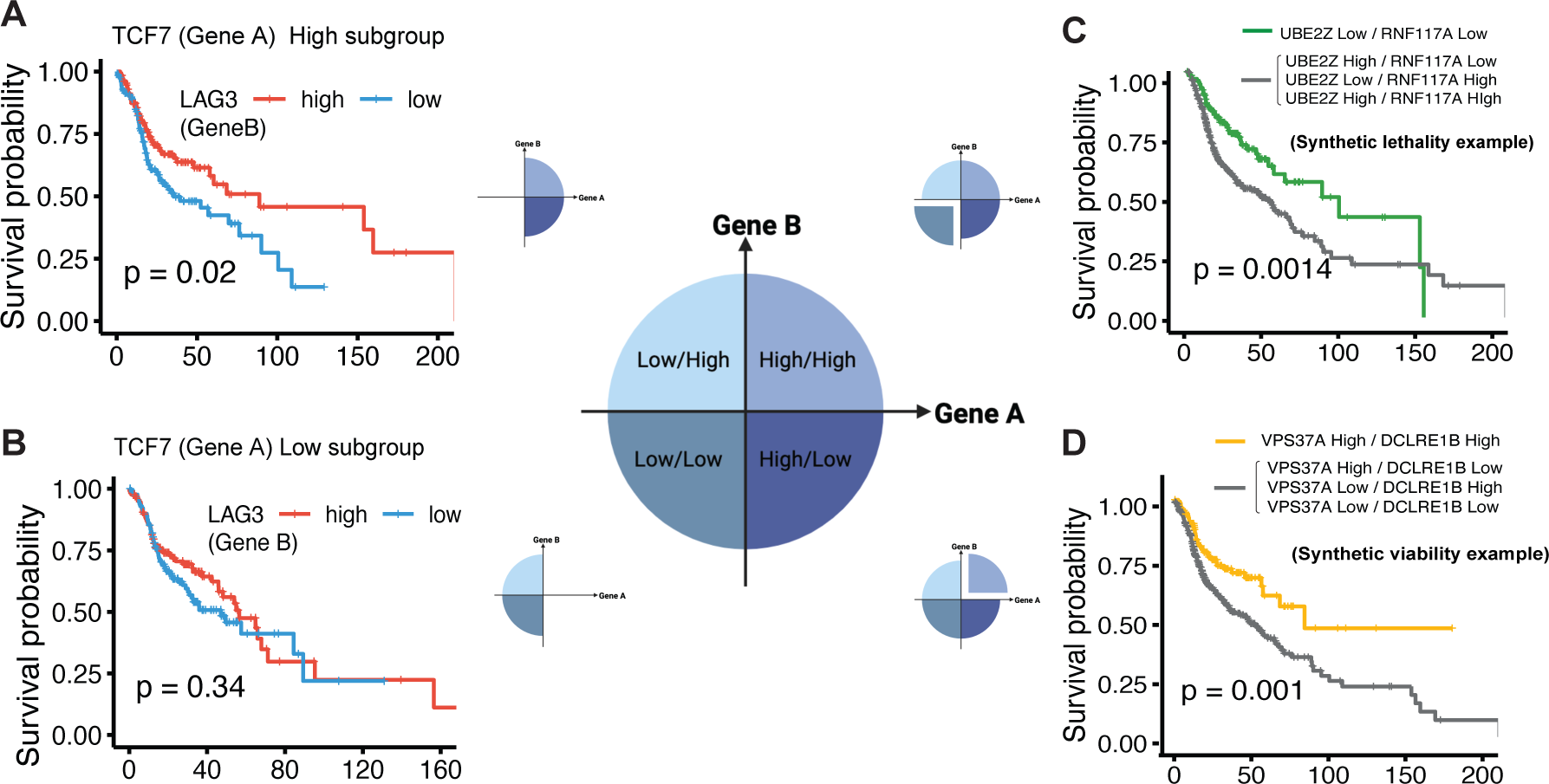
Prognostic relevance of gene-pair interactions as analyzed by CGPA. (A) Kaplan-Meier plot illustrating the survival impact of LAG3 levels in patients with high TCF7 expression, revealing a positive correlation between high LAG3 and improved survival, thereby indicating the prognostic significance of the TCF7-LAG3 interaction. (B) Kaplan-Meier plot for the TCF7-low group showing the lack of survival distinction based on varying LAG3 levels, suggesting a significant interaction effect of TCF7 and LAG3 on patient outcomes. (C) Visualization of synthetic viability with the VPS37A and DCLRE1B gene pair, where high expression levels of both genes correspond to better patient survival. (D) Representation of synthetic lethality in the context of UBE2Z and RNF117A gene pair, where low expression levels of both genes are associated with improved survival, implying a potential therapeutic strategy of dual-gene targeting.

### Gene network and subnetwork analysis

In this section, we further discuss the function of our gene network and subnetwork analysis implemented in CGPA. The function proves highly useful when investigating the functions of large gene network and gene sets with heterogenous functions^17^. Through gene set analysis, we can examine groups of genes that are collectively involved in influencing patient outcomes, which enables us to understand the broader biological context of individual gene signatures and aids in the validation of biomarker panels. Gene network analysis extends this approach by considering the interactions and regulatory relationships between genes within a network. It allows us to identify which genes are central ‘hubs’, potentially playing critical roles in tumorigenesis or immune-related functions, and to understand how disruptions in these networks may lead to disease progression. The current gene set analytical tools, such as GEIPIA2^18^ and SmulTCan^19^, lack the capability to examine subnetworks and their contribution within the prognostic model. To exemplify its utility in CGPA, we employ a composite gene signature, denoted as ISG.HY, derived from the tumor IFN–stimulated gene signature (ISG.RS) and genes associated with hypoxia. ISG.HY combines 38 genes from ISG.RS and 15 genes randomly selected from the hypoxia signature. In **Figure 5**, Panel A illustrates our application of Exploratory Graph Analysis (EGA) to dissect the ISG.HY gene set within the context of the TCGA Head and neck cancer dataset. EGA is an efficient subnetwork analysis tool, enabling us to better explore intra-gene relationships within complex pathways. This approach effectively segregates the ISG.HY gene set into three distinct subnetworks: ISG.HY1, ISG.HY2, and ISG.HY3. Remarkably, our analysis revealed that all hypoxia-related genes (such as VEGFA, ENO1 and MIF) were exclusively grouped in gene group 1 (ISG.HY.1). Other ISG.RS genes allocated to this group include HLA-G, HSD17B1, CA2, CCNA1, CXCL1, GALC, MCL1, ROBO1, SLC6A15, THBS1 and TIMP3, suggesting that these genes might be co-regulated or might interact in the hypoxic response. For example, HLA-G’s role in immune tolerance and MCL1’s involvement in apoptosis regulation could indicate mechanisms through which cells adapt to low oxygen environments. Genes associated with the interferon response and immune regulation were further stratified into group 2 and 3. When assessing the association of the ISG.HY signature with patient survival, we observed that overall gene set enrichment score (based on ssGSEA) did not exhibit significant correlations with patient survival. However, as depicted in the forest plot shown in Figure 5B, ISG.HY.1 emerged as a significant prognostic signature (hazard ratio of 1.88, p-value <0.001), while the other groups did not exhibit such significance. This finding suggests the potential role of ISG.HY.1 as a novel joint immune-hypoxia signature in the context of HNSC cancer progression and prognosis. In summary, the EGA function implemented in CGPA proves to be a useful tool that can aid in the refinement of existing prognostic gene signatures and thus enables the integration of both hypothesis-driven and data-driven approaches in signature discovery.

**Figure 5.**
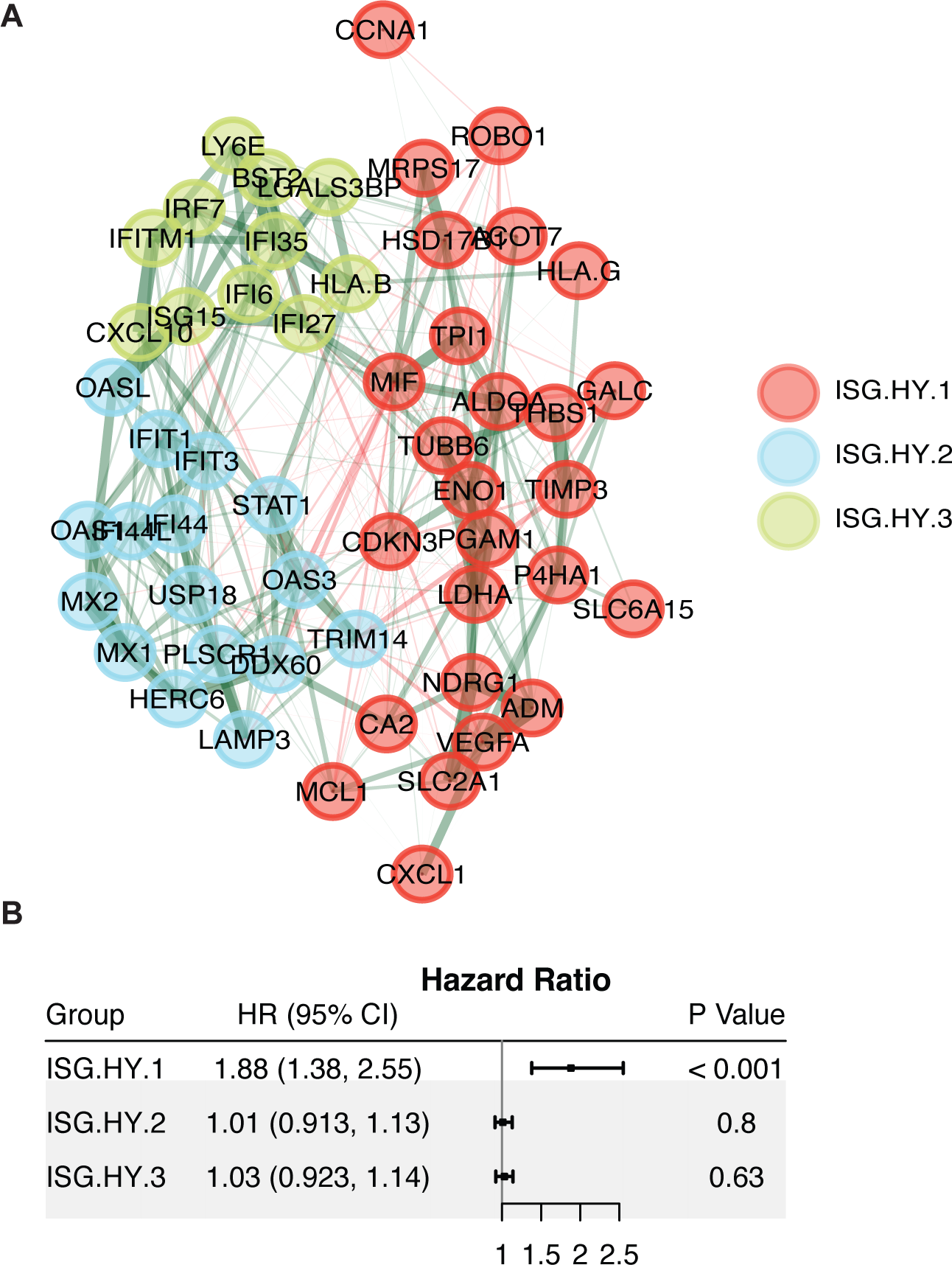
Dissection of the ISG.HY gene signature using CGPA’s Exploratory Graph Analysis (EGA) function. (A) The CGPA-EGA tool is applied to stratify the ISG.HY composite gene signature—consisting of 38 genes from the IFN-stimulated gene signature and 15 genes associated with hypoxia—into three functionally distinct subnetworks: ISG.HY1, ISG.HY2, and ISG.HY3, as demonstrated in the TCGA HNSC dataset. (B) The forest plot contrasts the prognostic value of each subnetwork, revealing ISG.HY1 as a significant prognostic indicator with a hazard ratio of 1.88 (p-value <0.001). This analysis underscores the capability of CGPA’s network and subnetwork analysis in understanding of complex gene interactions and refining prognostic gene signatures for improved cancer prognosis.

### Prognostic gene discovery in immunotherapy studies

In CGPA, we provide a designated page for discovering prognostic genes and gene modules based on immunotherapy datasets. We curated 33 published gene expression datasets comprising 2854 tumor samples from patients who received immune checkpoint inhibitor (ICI) therapy, with patient survival outcomes available. We offer all CGPA functions to facilitate comprehensive exploration and identification of potential prognostic biomarkers. Users can perform single-gene discovery to evaluate the prognostic significance of individual genes, conduct multivariable survival analysis to understand the combined impact of multiple genes on patient outcomes, discover gene pairs whose combined expression levels are predictive of survival, and analyze gene networks to identify key regulatory modules and pathways associated with immunotherapy response. This suite of tools allows researchers to thoroughly investigate the molecular mechanisms underlying treatment response and prognosis in the context of ICI therapy, providing valuable insights for the development and validation of personalized immunotherapy strategies.

## DISCUSSION

We introduced CGPA as an innovative online tool and interactive analysis portal aimed at addressing critical challenges in cancer genomics research. In comparison to existing tools such as GEPIA, GEPIA2^18^, and PRECOG^20^, CGPA offers a more comprehensive suite of advanced features, including the identification of UMOP genes, gene-pair analysis, and gene-set analysis, which are crucial for uncovering novel prognostic biomarkers and understanding complex molecular mechanisms in cancer progression. One of the key contributions of CGPA is its capability to customize survival models, enabling researchers to conduct Kaplan-Meier analysis with different cutoffs and Cox regression analysis with various clinical outcomes or cancer hallmarks. This flexibility in survival modeling is crucial for improving the accuracy and reliability of prognostic assessments and guiding personalized treatment strategies in oncology.

In current basic science literature, univariable comparisons based on Kaplan-Meier curves and the log-rank test are dominating methods to validate the prognostic values of a gene. While this approach is visually straightforward, it can lead to biased results due to the arbitrary cutoff of gene expression values to stratify patient groups. A prevalent misconception is that multivariable analysis, because it includes more variables in the model (thus using more degrees of freedom), can be less powerful than univariable analysis like KM and univariable Cox regression. Our analysis explicitly identified these missed opportunity genes (UMOP) genes, underscoring how significant insights can be overlooked in univariable analysis due to multiple reasons. Firstly, adjusting for confounding factors can reveal hidden associations between gene expression and survival outcomes, which might be diluted or distorted, such as by tumor purity. It is well known that gene expression from bulk tissue can be significantly influenced by tumor purity, and tumor purity, in some cases, can be a prognostic factor itself. Secondly, multivariable analysis often implicitly handles the interaction effects between genes and covariates, such as the CTL levels in our analysis. In the five immune-oncogenic cancer types we studied, it is well recognized that tumors classified as immune hot exhibit better prognosis than those classified as immune cold. However, survival differences highlighted by well-known immune genes like CD8A do not necessarily pinpoint the most crucial genes for biomarker discovery. This scenario is analogous to differential expression (DE) analysis when comparing tumor and normal tissues, where the most significantly differentially expressed genes are often keratins, yet these genes may not be the primary focus of interest. In our study across the five cancer types, the UMOP genes we discovered cover a diverse array of roles, including immune-related genes (HLA-C, HLA-H, HLA-B, and HLA-F), immune response genes (C1QB, C1QC, IL15, IFNB1, and SERPINA1), and immune modulators (TAP1 and GBP1). This diversity underscores the importance of multivariable analysis in uncovering critical genes that are missed in univariable analysis, thereby providing a more accurate and comprehensive understanding of the contribution of a targeted gene in cancer prognosis.

The TIDE^5^ method employs a similar Cox regression model focusing on the interaction between cytotoxic T lymphocytes (CTL) and specific genes, using z-scores from interaction terms to assess the effect of gene-CTL interactions on T cell dysfunction. Our gene-hallmark interaction analysis tool significantly expands upon TIDE by incorporating a wider array of clinical and molecular hallmarks, including top mutations, copy number alterations, androgen receptors, hypoxia, and various gene expression-based signatures. Essentially, TIDE could be considered a specialized instance within our broader model framework when focusing solely on the CTL hallmark. Our holistic gene-hallmark analysis is crucial for understanding the complex biology of cancer and developing targeted therapies. By analyzing gene-cancer hallmark interactions, our method not only identifies potential therapeutic targets but also aids in understanding how certain genes may influence or be influenced by these hallmarks. Moreover, even in the absence of significant interaction effects, multivariable analysis adjusting for cancer hallmarks can help identify prognostic genes that are most complementary to prognostic cancer hallmark signatures.

CGPA’s gene-pair analysis module offers valuable insights into the interplay between genes and their impact on patient survival. By integrating Kaplan-Meier plots and bivariable Cox regression analyses, CGPA enables researchers to identify synergistic or antagonistic gene interactions, shedding light on complex mechanisms underlying cancer progression. This functionality is particularly significant in uncovering synthetic lethality interactions, which may inform the development of targeted therapies and improve treatment outcomes for cancer patients. Additionally, CGPA’s gene-set analysis feature allows for the comprehensive evaluation of gene networks and pathways involved in cancer biology. Through single-sample Gene Set Enrichment Analysis (ssGSEA) and machine learning-based methods, CGPA facilitates the identification of gene subsets with significant prognostic value, enhancing our understanding of the molecular mechanisms driving cancer progression and guiding the development of novel therapeutic interventions.

Finally, CGPA provides two additional important features that set it apart from existing databases: (1) It provides a designated tab “ProgSplicing” to further explore the prognostic pan-cancer landscape of alternative splicing events, helping users to investigate genes with adverse prognostic effects across cancer types; (2) CGPA offers a dedicated portal for exploring prognostic gene modules using meticulously curated immunotherapy datasets. Altogether, the CGPA acts as a streamlined proactive and interactive gene-centric platform, greatly simplifying the task of prognostic biomarker research in oncology for clinicians and basic scientists. It is the first of its kind to provide multi-context insights from the cancer gene prognosis atlas, thereby bridging a critical gap in translational cancer research.

## METHODS

### Pan-cancer gene expression and survival data

In our study, we leveraged the Pan-Cancer normalized gene expression data from the PanCancer Atlas, accessible at https://gdc.cancer.gov/about-data/publications/pancanatlas. This dataset encompasses curated gene expression profiles across 33 cancer types, representing a total of 10,074 patients and covering 20,501 genes. In these 33 cancer types, eight cancers (LUAD, BRCA, HNSC, KIRC, LGG, LUSC, THCA, UCEC) have with over 500 samples. 10 other cancer types in the dataset have more than 200 samples, providing a comprehensive resource for the pan-cancer prognostic analysis. The survival outcomes used in this research is based on data from the TCGA Pan-Cancer Clinical Data Resource (TCGA-CDR). This resource offers a standardized dataset that incorporates curated survival endpoints (including OS, PFI, DFI, and DSS), along with endpoint usage recommendations tailored for each cancer type, enhancing the validity and rigorous of our findings. In our survival analysis, we prioritized primary tumor samples. Where multiple tumor samples existed for a single patient, we selected only one sample per patient for inclusion in the final survival analysis.

### Pan-cancer gene-based survival analysis

In this study, we conducted a pan-cancer gene-based survival analysis using univariable Cox modeling, focusing on overall survival (OS) and progression-free interval (PFI). The results were visually represented using forest plots, with cancer types ranked based on hazard ratios (HR). HR<1 indicated protective effects, while HR>1 suggested shorter survival times, and significance was determined by CI width at a 0.05 threshold. For cancer types that exhibited significant associations in the forest plots, we further explored the survival outcomes using Kaplan-Meier (KM) plots based on the optimal cutoff of gene expression values. The optimal cutoff was determined using the “surv_cutpoint” function implemented in the ‘survminer’ package. To investigate gene expression profiles across various cancer types and normal tissues, we used circular bar plots to rank cancer types based on normalized gene expression values. To gain insights into the biological and pathway context of the selected gene, a protein-protein interaction (PPI) network was constructed using the STRING API to contextualize gene function and pathways, limiting the number of retrieved interactions per gene to 10. These comprehensive analyses featured on the prognosis snapshot page of CGPA thus provide holistic insights into pan-cancer gene-based survival and gene expression patterns, aiding in our understanding of the relationships between genes, cancer types, and their potential implications in cancer biology and clinical outcomes.

### Single-gene multivariable analysis

To facilitate a more comprehensive single-gene multivariable analysis of gene-based survival, we provide a dedicated interactive survival analysis page equipped with highly customizable functions. This user-friendly page grants access to high-resolution KM plots generated under personalized modeling criteria. Below each KM plot, we provider the estimated Hazard Ratios (HR) and corresponding p-values obtained from the Cox proportional hazards modeling. In our analysis, we incorporated the capability for users to select specific covariates, including tumor purity, age, sex, and CTL (cytotoxic lymphocyte) levels, enabling them to conduct multivariable survival analyses tailored to their research objectives. To ensure consistent estimation of tumor purity, we downloaded a complete set of tumor purity data for all TCGA samples, which can be accessed at the following link: http://cistrome.org/TIMER/misc/AGPall.zip. It is worth noting that there were eight cancer types for which purity data was unavailable. Additionally, for seven sex-related cancer types, sex information was not integrated into the analysis, specifically CESC, OV, PRAD, TGCT, UCEC, UCS, and BRCA. In the multivariable analysis tab, we also compiled a pre-calculated list of the top prognostic genes for each cancer type, encompassing both mRNA and long non-coding RNA (lncRNA) categories. It is important to mention that we kept the mRNA and lncRNA datasets separate, as they underwent distinct preprocessing pipelines. This approach ensures that users have access to the most comprehensive and personalized gene-based survival information for their specific research needs.

### Gene hallmark analysis

In our gene hallmark analysis, we considered age, sex, the top mutated genes, copy number variations (CNV) specific to each cancer type, tumor mutational burden (TMB), immune scores (including CYT and CTL), and the androgen receptor (AR) as cancer hallmarks in the prognostic models. We explored two models to test the interaction between these hallmarks and the selected gene: the GHI full model (Hazard ∼ a.Gene + b. Hallmark + c.Gene x Hallmark) and the GHI partial model (Hazard ∼ a.Gene + c.Gene x Hallmark). The visualization of the results was accomplished through circos plots, highlighting significant interaction connections at a significance level of 0.05. All gene set signature (e.g. CYT and CTL) analysis were calculated based on mean values or single-sample gene set enrichment analysis (ssGSEA). All analyses offered the option to adjust for tumor purity, ensuring a comprehensive examination of the interplay between hallmarks, genes, and their impact on survival outcomes across diverse cancer scenarios.

### CGPA two-gene analysis

The CGPA toolkit introduces a comprehensive module dedicated to exploring the prognostic implications of gene-pair interactions in cancer. This analysis leverages a dual-method approach, incorporating both Kaplan-Meier (KM) plots and bivariable Cox regression analyses, to evaluate the survival impact associated with the interaction of two genes. The KM plots are employed as a fundamental visual tool within the CGPA toolkit, enabling the intuitive representation of survival probabilities over time for patients based on the interaction between pairs of genes. By dichotomizing gene expression into high and low categories, researchers can discern the prognostic value through a clear observation of survival outcomes among these defined groups. Should an interaction effect exist, the survival curves will be further separated within patient subgroups stratified by the expression of one gene, as exemplified in the case genes presented in Figure 3. Importantly, enabling different cutoffs of KM plots in patient stratification allows for a more thorough examination of interaction effects in a non-linear and discontinuous manner. The analysis extends to explore phenomena such as synthetic viability and lethality through gene-pair interactions. Synthetic viability, as observed with the VPS37A and DCLRE1B gene pair, indicates scenarios where high expression levels of both genes correlate with improved survival outcomes. Conversely, synthetic lethality is demonstrated through the UBE2Z and RNF117A gene pair, where low expression of both genes suggests a favorable prognostic outcome, potentially guiding the development of targeted therapies.

Complementing the KM plots, bivariable Cox regression analyses offer a more formal statistical examination by including interaction terms between the two genes of interest. This enables the assessment of not only the individual but also the combined effects of gene expressions on survival. The CGPA two-gene analysis also incorporates an innovative feature to examine the prognostic value of the ratio of expression values between two genes, offering a notable advantage by identifying interaction relationships that may be missed by the additive models. This ratio also serves as a robust prognostic signature, especially useful in cases of high correlation between genes. Moreover, it provides an additional normalization, using one gene’s expression as self-control for the other, thus mitigating batch effects and biases in gene expression normalization methods.

### Multi-Gene Prognostic Analysis

The CGPA introduces a module for multi-gene analysis, designed to dissect complex gene networks and sets with heterogeneous functions. To evaluate the prognostic significance of gene sets, we leverage single-sample Gene Set Enrichment Analysis (ssGSEA), employing it alongside the median as methods for calculating combined scores within survival analysis. The analysis framework allows for the utilization of both univariable Cox regression analysis and machine learning-based methods, specifically the gradient boosting method^21^, to identify top prognostic genes within the input set. This dual approach ensures a comprehensive evaluation of gene significance, incorporating both traditional statistical models and advanced predictive algorithms to highlight genes with the most substantial impact on prognosis. The exploration of gene networks and subnetworks is conducted using EGAnet^22^, a gene community-based approach that facilitates the understanding of complex intra-gene relationships. This method is able to classify genes into distinct subnetworks based on their interaction patterns, thereby elucidating potential co-regulation or interaction mechanisms among genes within specific pathways or responses, such as hypoxia or immune regulation. To further explore the patterns of gene interactions, the module incorporates tools for inter-gene correlation analysis based on igraph, as well as principal component analysis (PCA) for visualization purposes.

These tools allow users to visually assess the correlation patterns among genes within the network, aiding in the selection or exclusion of genes for the panel construction. By integrating these methods, the multi-gene analysis function of the CGPA provides a seamless blend of hypothesis-driven and data-driven approaches, fostering the discovery and validation of meaningful gene signatures in cancer prognosis.

## Data and code availability

CGPA is publicly available as a web-based interactive tool at cgpa.moffitt.org. This platform provides researchers with easy access to a comprehensive suite of tools for prognostic gene discovery and analysis in immunotherapy studies. The source code and associated R packages used in the development of CGPA have been deposited in a GitHub repository. Researchers can access these resources, along with detailed instructions for installation and use, at https://github.com/wanglab1/CGPA. This repository ensures transparency and reproducibility of the methods used in CGPA, allowing users to customize and extend the functionalities as needed for their specific research purposes. Additionally, all data included in the CGPA web tool, including both PanCancer TCGA datasets and immunotherapy datasets, is available for download at https://github.com/wanglab1/CGPA/tree/main/PanCancer_data and https://github.com/wanglab1/CGPA/tree/main/IOtherapy_data. The TCGA CDR outcome data (recommended for survival analysis) were downloaded from the “TCGA-CDR-SupplementalTableS1.xlsx” file available on the PanCanAtlas website: https://gdc.cancer.gov/about-data/publications/pancanatlas. The alternative splicing datasets generated by SpliceSeq and SpIAdder were acquired from oncosplicing.com, with supplementary data from all cancer types (not available on the website) provided by the OncoSplicing author team. The ICI datasets preprocessed and analyzed by this study are available from the Gene Expression Omnibus (GEO) repository under the accession numbers: GSE111636, GSE248167, GSE173839, GSE194040, GSE241876, GSE179730, GSE162137, GSE165252, GSE159067, GSE195832, GSE67501, GSE135222, GSE221733, GSE126044, GSE100797, GSE115821, GSE131521, GSE78220, GSE91061, GSE96619, GSE202687. We also included pre-analyzed gene expression data from large ICI projects, including IMvigor210 (based on the R package “IMvigor210CoreBiologies”), harmonized KIRC datasets from the supplementary files of publication PMID: 32472114, Javeline101 from the supplementary files of publication PMID: 32895571, harmonized SKCM datasets (available at https://github.com/ParkerICI/MORRISON-1-public), and Kallisto data from https://github.com/hammerlab/multi-omic-urothelial-anti-pdl1/tree/master.

## Author contributions

BW, XY, GG, ARM, SL, TL, PJ, and XW performed research. BC, GG, and ARM built the CGPA interactive framework. JRC provided critical expertise in cancer biomarker research and edited the manuscript. PJ and XW conceptualized the study. XW supervised the research and wrote the manuscript.

## Acknowledgments

This work has been supported in part by a National Institutes of Health grant [R01DE030493 to X.W], the State of Florida Bankhead-Coley Cancer Research Program, infrastructure research grant 23B16, and Biostatistics and Bioinformatics Shared Resources at the H. Lee Moffitt Cancer Center & Research Institute, an NCI-designed Comprehensive Cancer Center (P30-CA076292).

